# Design of a de novo aggregating antimicrobial peptide and bacterial conjugation delivery system

**DOI:** 10.1101/325621

**Authors:** Logan T. Collins, Peter B. Otoupal, Colleen M. Courtney, Anushree Chatterjee

## Abstract

Traditional antibiotics are reaching obsolescence as a consequence of antibiotic resistance; therefore novel antibiotic approaches are needed. A recent non-traditional approach involves formation of protein aggregates as antimicrobials to disrupt bacterial homeostasis. Previous work on protein aggregates has focused on genome mining for aggregation-prone sequences in bacterial genomes rather than on rational design of aggregating antimicrobial peptides. Here, we use a synthetic biology approach to design an artificial gene encoding the first de novo aggregating antimicrobial peptide. This artificial gene, *opaL* (overexpressed protein aggregator Lipophilic), disrupts bacterial homeostasis by expressing extremely hydrophobic peptides. When this hydrophobic sequence is disrupted by acidic residues, consequent aggregation and antimicrobial effect decreases. Further, to deliver this artificial gene, we developed a probiotic approach using RK2, a broad host range conjugative plasmid, to transfer *opaL* from donor to recipient bacteria. We utilize RK2 to mobilize a shuttle plasmid carrying the *opaL* gene by adding the RK2 origin of transfer. We show that *opaL* is non-toxic to the donor, allowing for maintenance and transfer since its expression is under control of a promoter with a recipient-specific T7 RNA polymerase. Upon mating of donor and recipient *Escherichia coli*, we observe selective growth repression in T7 polymerase expressing recipients. This technique could be used to target desired pathogens by selecting pathogen-specific promoters to control *opaL* expression. This system provides a basis for the design and delivery of novel antimicrobial peptides.

**Importance:** The growing threat of antibiotic resistance necessitates new treatment options for bacterial infections that are recalcitrant to traditional antimicrobials. Existing methods usually involve small-molecule compounds which interfere with essential processes in bacterial cells. By contrast, protein aggregates operate by causing widespread disruption of bacterial homeostasis and may provide a new method for combating infections. We used rational design to create and test an aggregating de novo antimicrobial peptide, OpaL. In addition, we employed bacterial conjugation to deliver the *opaL* gene from donor bacteria to recipient bacteria while using a strain-specific promoter to ensure that OpaL was only expressed in targeted recipients. To the best of our knowledge, this represents the first design for a de novo peptide with aggregation-mediated antimicrobial activity. We envision that OpaL’s design parameters could be used in developing a new class of antimicrobial peptides to help treat antibiotic resistant infections.

## 1 Introduction

Traditional antibiotic treatments are losing efficacy as bacteria gain resistance to available antimicrobial compounds. This resistance arises through a variety of mechanisms such as altered target sites, exclusion from the cell, enzymatic degradation or modification of antibacterial compounds, and efflux pumps (1). Carbapenem resistant enterobacteriaceae have been reported in several nations including the United States, India, the UK, and others (2). These bacteria often cause infections which are untreatable with most or even all current antibiotics. We may be approaching a post-antibiotic era in which antibiotic treatments are no longer functional (3). With such prospects, new treatments are needed to protect public health.

The advent of synthetic biology has provided tools to develop improved antimicrobial strategies. Engineered biofilm-disrupting bacteriophages reduce biofilm mediated resistance. (4). Other modified bacteriophages deliver CRISPR systems to inactivate antibiotic resistance genes in pathogenic bacteria. In addition, CRISPR libraries have been used to study how antibiotic resistance evolves by manipulating gene expression under antibiotic stress conditions (5, 6). Other examples include rearrangement of domains within polyketide synthase complexes to enable the production of new polyketide antimicrobials (7). Recently, probiotic bacteria have been engineered to synthesize autoinducers that inhibit virulence genes in infectious microorganisms (8). Other recombinant probiotic bacteria express Shiga toxin receptors that neutralize some of the virulence factors produced by pathogenic *E. coli* (9). Here we use synthetic biology principles to develop a novel de novo aggregating antimicrobial peptide that can be used to kill bacteria. We also repurpose an RK2-mediated bacterial conjugation system to develop a probiotic approach for delivering this novel peptide.

Antimicrobial peptides (AMPs) have shown promise as antibacterial treatments. AMPs, many of which contain hydrophobic and cationic domains (10), are expressed naturally by numerous organisms to combat bacterial proliferation. These AMPs bind to bacterial cells by ionic interactions with negatively charged membrane components before inserting into the phospholipid bilayer and disrupting membrane integrity by forming pores. Alternatively, some AMPs may enter the cell and disrupt specific intracellular processes such as protein synthesis and DNA replication (10). Though AMPs are a relatively new area, a few like polymyxin B and colistin, are established clinical agents (11). Numerous AMPs have shown potential in pre-clinical contexts and are undergoing development. (10, 12). Using principles derived from naturally occurring AMPs, synthetic biology has enabled the construction of enhanced AMPs.

For instance, consensus sequences have been derived from libraries of naturally occurring amphipathic AMPs to optimize pore formation (13). Another method involves screening AMP libraries and linking the most effective peptides to maximize their potency (14). In addition, rational sequence modifications to adjust properties such as helicity and charge, incorporation of unnatural amino acids, and chemical modifications have been used to improve the activity of preexisting AMPs (10).

Here we investigate a new category of AMPs that operate via an aggregative mechanism. The toxicity of aggregating peptides arises from mechanisms including the disruptive interaction of exposed hydrophobic side chains with cellular proteins (15), induction of oxidative stress, overload of proteolytic machinery, disruptive interaction with membranes, and co-aggregation with endogenous macromolecules (16). When in their aggregate-prone states, the peptides associated with diseases often exhibit β-sheet-rich structures (17). Mechanistically, β-sheets and hydrophobicity have been supported by thermodynamic models as driving protein aggregation (18). Aggregating peptides have been observed in a number of human pathologies including Alzheimer’s disease, prion diseases, Parkinson’s disease, Huntington’s disease, and sickle cell anemia (19).

Using aggregating AMPs as antimicrobials may provide a novel strategy to counter antibiotic resistance. Aggregating AMPs utilize an unexploited mechanism of antimicrobial toxicity wherein instead of binding a particular target site on a bacterial macromolecule, these can cause widespread disruption of homeostasis in bacteria. This may prevent resistance from emerging since many resistance phenotypes involve target site alterations. Recently, Bednarska et al. demonstrated the potential of using aggregating peptides as antibiotics (20). The authors investigated the antimicrobial activity of short peptides derived from bacterial aggregate-promoting sequences in *Staphylococcus epidermidis* proteome by computationally predicting aggregation propensity using a statistical thermodynamics algorithm called TANGO (18). The resulting peptides were constructed by solid-phase peptide synthesis and exogenously delivered to bacterial cells. A small fraction of the screened peptides showed significant therapeutic activity which highlights the promise of aggregating peptides as antimicrobials. However, a major limitation of this approach was that these peptides were not rationally designed and involved screening hundreds of candidate molecules with limited success.

Here we rationally design the first (to the best of our knowledge) de novo aggregating antimicrobial peptide, OpaL (Overexpressed protein aggregator Lipophilic), by choosing numerous hydrophobic amino acid residues to maximize protein aggregation rather than mining inclusion-forming sequences from bacterial genomes (Fig. 1A). The *opaL* gene was specifically designed to disrupt bacterial homeostasis by overexpressing hydrophobic, aggregate-prone peptides. Using computational prediction and confirmation by experimental methods, we show that OpaL causes aggregation in bacteria leading to bacterial lethality. A similar control synthetic peptide which includes numerous acidic residues exhibits significantly less toxicity. We use the broad host range conjugative plasmid RK2 to mobilize a pET11a-based shuttle plasmid which carries *opaL* to transfer the antimicrobial peptide from donor to recipient bacteria. We show that this conjugative system can successfully target the strain of interest using a strain-specific promoter. Our work provides a new therapeutic strategy which may have applications in the clinical setting.

**Figure 1.**
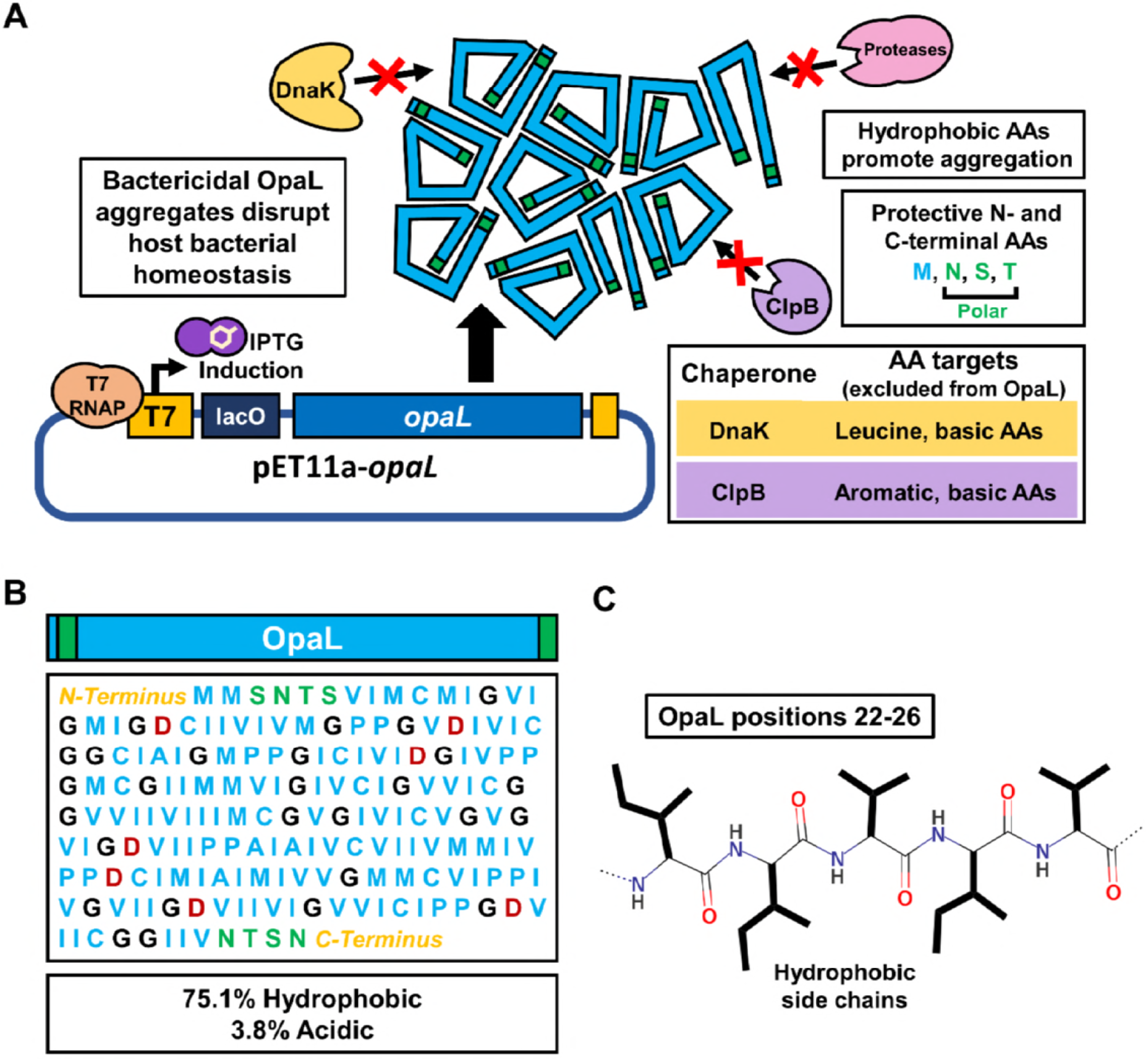
Rational design of antimicrobial hydrophobic peptide. **(A)** The pET11a-*opaL* vector expresses OpaL from the strong T7 promoter. Using T7 restricts OpaL expression to the target strain, *E. coli* BL21 (DE3). OpaL forms hydrophobic aggregates, disrupts bacterial homeostasis, and kills the host microorganism. Polar terminal patches were incorporated to increase stability by disfavoring interactions with bacterial proteases (25). Leucine, aromatic amino acids, and basic amino acids were excluded from the OpaL sequence because these residues tend to favor interactions with DnaK (26) and the disaggregase ClpB (27). Numerous (76.3%) hydrophobic residues facilitate OpaL’s ability to form toxic intracellular aggregates (15, 16). **(B)** The OpaL peptide’s amino acid sequence is primarily hydrophobic (blue). Exceptions include its polar N- and C-terminal patches (green) for stability, scattered glycines (black) for conformational flexibility, and a small number of aspartic acid residues (red) to further disfavor binding by the chaperone DnaK (28). **(C)** Structure of OpaL residues 22-26. OpaL’s bulky nonpolar side chains create a hydrophobic chemical environment.

## 2 Materials and methods

### 2.1 Strains, plasmids, kits, and gene synthesis

The pET11a-*opaL* and pET11a-*opaLacidic* vector designs were constructed by GenScript using their artificial gene synthesis and custom cloning services. The *opaL* open reading frame, *opaLacidic* open reading frame, RK2 OriT, and chloramphenicol resistance (CmR) gene were artificially synthesized. RK2 was obtained in *E. coli* C600 (ATCC^®^ 37125^TM^). We used a *Mix & Go E. coli* Transformation Kit from Zymo Research to induce chemical competence in the *E. coli* C600 (RK2) before transforming with pET11a-*opaL*. To prevent loss of RK2, we grew these cells under kanamycin selection. *E. coli* C600 (RK2, pET11a-*opaL*) and *E. coli* C600 (RK2) were subsequently used as donor bacteria.

The pHL662 plasmid, carried by *E. coli* XL1 Blue, was Addgene vector 37636 (21). The *E. coli* XL1 Blue was used as recipient bacteria for measuring the mating frequency. We isolated pHL662 using a Zymo Research Zyppy Plasmid Miniprep Kit. Chemically competent *E. coli* BL21 (DE3) were acquired from NEB and transformed with pHL662. These *E. coli* BL21 (DE3) (pHL662) were used as recipients in the mating-toxicity assays. Separate samples of *E. coli* BL21 (DE3) were also transformed with pET11a-*opaL* for toxicity assays. *E. coli* NEB10-β was acquired from NEB and used as a host for propagating the pET11a-OriT vector. The pUV145 plasmid (22), was carried in an *E. coli* DH5α host. *E. coli* DH5α carrying pUV145 were employed in our three strain mating-toxicity assay. A list of all strains and plasmids used for this work is presented in supplementary table S1.

### 2.2 Culture conditions

Growth media included Luria Bertani (LB) broth (liquid medium) and LB agar (solid medium) with selective antibiotics as described for each experiment. Liquid cultures were incubated at 37°C in an orbital shaker or in a Tecan GENios plate reader. When using the plate reader, cultures were set to shake for 10 minutes, stand idle for 10 minutes, and then shake for an additional 10 seconds prior to taking a measurement. Solid cultures were grown in a stationary incubator at 37°C. The *opaL* gene and the *opaLacidic* gene were induced using IPTG at concentrations of 1.0 or 0.1 mM as described for each experiment. GFP from pHL662 and mCherry from pUV145 were induced using IPTG at concentrations of 1.0 mM. Except where otherwise noted, ampicillin was used to maintain pET11a-*opaL* and pET11a-*opaLacidic* and kanamycin was used to maintain RK2, pHL662, and pUV145.

### 2.3 Molecular cloning

The pET11a-Δ*opaL* control plasmid was prepared by removing *opaL*’s open reading frame (positions 6124 to 6690). To accomplish this, pET11a-*opaL* was first propagated in *E. coli* DH5α to facilitate DNA methylation. Primers were designed to amplify the part of pET11a-*opaL* which excludes the opaL-containing sequence between pET11a-*opaL* plasmid’s NdeI and BamHI cut sites. The forward primer (5’-ggaaggggatccggctgctaacaaag-3’) still retained the original BamHI cut site sequence, while the reverse primer (5’-gaggagggatcctatatctccttcttaaagttaaacaaaat-3’) included a small overhang which replaced the NdeI cut site with another BamHI cut site upon amplification. After amplifying this sequence, the PCR product was double digested with DpnI (a methylation-dependent restriction enzyme) and BamHI. The purpose of using DpnI was to degrade any remaining background DNA which still contained *opaL.* Next, the linear vector was ligated overnight and then electroporated into *E. coli* NEB10-β. A transformant was picked and grown in liquid media overnight. The pET11a-OriT plasmid was miniprepped from this culture and then confirmed to have the correct size by performing gel electrophoresis alongside a sample of pET11a-*opaL*. Maps of plasmids used for this work are presented in supplementary Fig. S1.

### 2.4 Toxicity assay using growth curves

Overnight cultures (three biological replicates) of *E. coli* BL21 (DE3) carrying pET11a-*opaL, E. coli* BL21 (DE3) carrying pET11a-*opaLacidic*, *E. coli* BL21 (DE3) carrying pET11a-Δ*opaL*, and *E. coli* BL21 (DE3) without any plasmids were diluted 1×10^−4^ and incubated for 2 hours. Samples from each culture were diluted 1:50 into fresh media with 1.0 mM IPTG, 0.1 mM IPTG, and 0.0 mM IPTG in a 96 well plate. Absorbance values were measured every 20 minutes for 20 h using the Tecan Genios plate reader settings described earlier. These data were normalized by subtracting the absorbance of the media and dividing by the OD at t=0 for each sample.

### 2.5 Toxicity assay using Colony Forming Units (CFUs)

We tested *opaL*’s antibacterial activity in *E. coli* BL21 (DE3) carrying pET11a-*opaL* and *E. coli* BL21 (DE3) carrying *pET11a-opaLacidic*. Overnight starter cultures (three biological replicates) were diluted 1:1000 and incubated for 2 hours. Serial dilutions of these exponential cultures were plated on solid medium to obtain CFUs at t=0 h. The cultures were further diluted 1:100. Immediately after these dilutions, we split the cultures into control and experimental tubes and then added IPTG to the experimental tubes (1.0 mM final concentration) in order to induce *opaL* expression. Serial dilutions were then plated on solid media at t=2 h and t=4 h.

### 2.6 Computational prediction of aggregation using TANGO

The online TANGO platform (http://tango.crg.es/protected/academic/calculation.jsp) was used to predict percent aggregation for OpaL and OpaLacidic. The same parameters were used for both peptides. The peptide concentrations were set at 2.0 mM, the ionic strengths at 0.025 M, the pH at 7.5, and the temperature at 37°C. The N- and C-termini were given the default parameter of not having any chemical modifications.

### 2.7 Computational structure prediction using QUARK

The online QUARK platform (23) (https://zhanglab.ccmb.med.umich.edu/QUARK/) was used to predict tertiary structures for OpaL and OpaLacidic. The raw amino acid sequences were inputted into the algorithm and the results retrieved in PDB file format. From these files, 3D graphics were created with DeepView v4.1.0 (24). DeepView was also used to compute OpaL and OpaLacidic’s solvent-exposed hydrophobic surface areas.

### 2.8 Nile red aggregation assay

Overnight cultures (ten biological replicates) of *E. coli* BL21 (DE3) carrying pET11a-*opaL, E. coli* BL21 (DE3) carrying pET11a-*opaLacidic*, *E. coli* BL21 (DE3) carrying pET11a-OriT, and *E. coli* BL21 (DE3) without any plasmids were diluted 1:10 into 900 μL of fresh media and incubated for 1 h before being pelleted, washed, and resuspended in Phosphate Buffered Saline (PBS). Nile red was added to a final concentration of 1.0 μg/mL. Absorbance was measured at 590 nm and fluorescence was measured at 590 nm excitation and 610 nm emission in a Tecan Genios plate reader. Next, IPTG was added to five of the replicates to a concentration of 1.0 mM and the other five replicates to a concentration of 0.1 mM. These samples were then incubated at 37°C for 2 h with shaking. Absorbance and fluorescence were measured again at the same wavelengths. Initial and final fluorescence values were normalized to the corresponding absorbance values and the overall normalized changes in fluorescence were calculated.

### 2.9 Mating assay

Mating assays were performed to confirm that RK2 and pET11a-*opaL* are capable of conjugative transfer. 1 mL overnight cultures (three biological replicates) of *E. coli* C600 donors carrying both RK2 and pET11a-*opaL*, *E. coli* C600 donors carrying the only RK2, and *E. coli* XL1 Blue recipients carrying pHL662 were pelleted and washed to remove antibiotics before being resuspended in 250 μL of media. The volumes of these cultures were adjusted to have equal OD values before mating cultures with 1:5 donor to recipient ratios were made by volume. *E. coli* C600 (RK2, pET11a-*opaL*) and *E. coli* C600 (RK2) were each separately paired with the recipients. 20 μL of the mating cultures were spotted on LB agar plates without selection and incubated for 5 hours. Next, we cut out solid agar slices with the spots and transferred them to liquid cultures without selection. After 1 h of incubation, the cultures were diluted 1:10,000 and plated on X-Gal with appropriate antibiotics.

X-Gal allowed distinction between blue donor (*E. coli* C600) colonies and white transconjugant (*E. coli* XL1 Blue) colonies. *E. coli* XL1 Blue possess the Δ*lacZ* genotype and so cannot metabolize X-Gal to produce blue pigment. Donors and transconjugants with both RK2 and pET11a-*opaL* were selected with chloramphenicol while donors and transconjugants with only RK2 were selected with ampicillin. Control X-Gal plates without antibiotics were made for each mating culture. Mating frequencies were determined by taking the ratio of transconjugant colonies over total recipient colonies.

### 2.10 Two strain mating-toxicity assay using CFUs

Mating-toxicity assays using CFUs demonstrated the functionality of our bacterial conjugation delivery system for transferring *opaL* to target bacteria. 1 mL overnight cultures were made with three biological replicates of *E. coli* C600 donors carrying both RK2 and pET11a-*opaL*, *E. coli* C600 donors carrying only RK2, and *E. coli* BL21 (DE3) recipients carrying pHL662. These cultures were then diluted 1:100 in 5 mL LB medium and incubated for 3 h, followed by pelleting and washing twice with LB to remove antibiotics, and then resuspension in 250 μL of fresh medium. Culture volumes were adjusted to give approximately equivalent OD values. Next, donor and recipient strains were mixed to create mating cultures with 1:3 donor to recipient ratios. Mating cultures were spotted onto 1.0 mM IPTG plates without selection and incubated for 5 h. We cut out the solid agar slices with mating spots, transferred them each into 1 mL of PBS, and vortexed thoroughly to resuspend the mated bacteria. These cells were diluted to 1×10^−5^ and plated on LB agar plates containing kanamycin (to maintain pHL662) and 1.0 mM IPTG. GFP-expressing (recipient) colonies and non-fluorescent (donor) colonies were counted using 470 nm excitation and 530 nm emission wavelengths.

### 2.11 Two strain mating-toxicity assay using fluorescence growth curves

Mating-toxicity assays using fluorescence growth curves further showed the functionality of our bacterial conjugation delivery system for the *opaL* gene. 1 mL overnight cultures were made with four biological replicates of *E. coli* C600 donors carrying both RK2 and pET11a-*opaL*, *E. coli* C600 donors carrying only RK2, and *E. coli* BL21 (DE3) recipients carrying pHL662. The overnight cultures were washed twice to remove antibiotics, and resuspended in 250 μL of LB media. Culture volumes were adjusted to give approximately equivalent OD values. Next, donor and recipient strains were mixed to create mating cultures with 1:1 donor to recipient ratios. Mating cultures were spotted onto 1.0 mM IPTG plates without selection and incubated for 5 h. We cut out the solid agar slices with mating spots and transferred them to liquid media, where they were incubated for 1 h. The cultures were diluted 1:20 into fresh media with 1.0 mM IPTG in a 96 well plate. GFP fluorescence was measured with 485 nm excitation and 535 nm emission every 20 minutes for 20 h using the Tecan Genios plate reader settings described earlier. These data were normalized with the media’s autofluorescence values.

### 2.12 Three strain mating-toxicity assay using CFUs

We performed a three strain mating-toxicity assay to demonstrate that this conjugation based delivery approach functions effectively when bacteria other than the donors and the targeted recipients are present. 1 mL overnight cultures were made with three biological replicates of *E. coli* C600 donors carrying both RK2 and pET11a-*opaL*, *E. coli* C600 donors carrying only RK2, *E. coli* BL21 (DE3) recipients carrying pHL662, and *E. coli* DH5α recipients carrying pUV145. As in the two strain assay, the overnight cultures were diluted 1:100 in 5 mL LB and incubated for 3 h, followed by washing twice to remove antibiotics, and resuspension in 250 μL of media. Culture volumes were adjusted to give approximately equivalent OD values. Donor *E.coli* C600, recipient *E. coli* BL21 (DE3), and recipient *E. coli* DH5α were mixed to create mating cultures with ratios of 1:1:2 respectively. Mating cultures were spotted onto 1.0 mM IPTG plates without selection and incubated for 5 hours. Equivalent volumes of *E. coli* DH5α alone were spotted onto 1.0 mM IPTG plates without selection and incubated for 5 h. We cut out the solid agar slices with mating spots, transferred them each into 1 mL of PBS, and vortexed thoroughly to resuspend the mated bacteria. These cells were diluted to 1×10^−5^ and plated on kanamycin (to maintain pHL662 and pUV145) and 1.0 mM IPTG. GFP-expressing (target recipient) colonies and non-fluorescent (donor) colonies were counted using 470 nm excitation and 530 nm emission wavelengths, while mCherry-expressing (non-target recipient) colonies were counted using 540 nm excitation and 590 nm emission wavelengths.

## 3 Results

### 3.1 Design of de novo aggregating antimicrobial peptide

To design the OpaL peptide’s amino acid sequence, we considered (i) potential for aggregation, (ii) potential for molecular chaperones and proteases to mitigate the toxic effects of aggregation, and (iii) the polypeptide’s half-life in the cell (Fig. 1A). To promote insolubility in the aqueous cytosolic environment, 139 amino acids of the chosen 185 residue sequence (75.1%) of OpaL possess hydrophobic side chains. These residues include 4 (2.2%) alanine, 15 (8.1%) cysteine, 53 (28.6%) isoleucine, 17 (9.2%) methionine, 14 (7.6%) proline, and 36 (19.5%) valine (Fig. 1B-C). Because molecular chaperones can sequester improperly folded proteins for degradation, several molecular chaperone-targeted amino acids and sequence motifs were avoided in the design of the polypeptide (25). These include aromatic and basic residues (which favor binding by the disaggregase ClpB) (26) as well as leucine and hydrophobic motifs flanked by basic residues (which are targeted by the chaperone DnaK) (Fig. 1A) (27). Since DnaK binding also disfavors acidic amino acids (28), seven aspartic acid residues were incorporated into the highly hydrophobic OpaL sequence. These aspartic acids were all spaced at least twelve residues apart from each other to minimally interfere with hydrophobicity-mediated aggregation. Since protein half-life is strongly influenced by terminal amino acid sequences, several protective polar amino acids (serine, asparagine, and threonine) were incorporated to avoid degradation by proteases which vastly shorten half-lives by recognizing bulky terminal hydrophobic residues (25). N-formylmethionine was still allowed at the N-terminus since, although it is hydrophobic, N-formylmethionine promotes long protein half-lives (29). Since the N-formylmethionine found in bacterial proteins is often cleaved by intracellular proteases, the next several chosen residues were also among those that promote long half-lives. These design parameters may also provide a general template for future variations of the OpaL peptide.

To verify that OpaL’s antimicrobial activity arose mainly from its highly hydrophobic character (75.1% hydrophobic amino acids), we created a similar 184 residue control peptide, OpaLacidic, with greatly increased content of acidic amino acids. This peptide includes 30 aspartic acid residues comprising 16.3% of the sequence (supplementary Fig. S3). To facilitate charge-charge repulsion and interfere with hydrophobic aggregation (from OpaLacidic’s 70.7% hydrophobic residues), these aspartic acids were placed every five residues over the majority of the sequence.

### 3.2 Impacts of OpaL on bacterial growth

We demonstrated OpaL’s toxic effect by expressing it under the control of the strong T7 promoter on the medium-copy pET11a-*opaL* shuttle plasmid in *E. coli* BL21 (DE3). In pET11a derived vectors, a lac operator is located directly downstream of the T7 promoter, allowing induction with isopropyl β-D-1-thiogalactopyranoside (IPTG). When measuring the growth of bacterial cultures of BL21 (DE3) carrying pET11a-*opaL* using optical density (OD), we found that inducing OpaL expression with 1.0 mM IPTG completely precluded growth (Fig. 2A). When using 0.1 mM IPTG instead, OpaL expression still decreased mean OD (Fig. 2B), though some growth occurred. In absence of IPTG, BL21 (DE3) carrying pET11a-*opaL* showed a longer lag time compared to *E. coli* BL21 (DE3) not carrying any plasmids, potentially due to leaky expression from the *opaL* gene (Fig. 2C). In contrast, *E. coli* BL21 (DE3) expressing OpaLacidic displayed less toxicity than OpaL, albeit with lengthened lag times relative to control bacteria not carrying any plasmids (Fig. 2A-C). To further investigate this, we deleted the open reading frame of *opaL* to create the control plasmid pET11a-Δ*opaL*. *E. coli* BL21 (DE3) carrying pET11a-Δ*opaL* demonstrated significantly higher growth than cells expressing OpaL or OpaLacidic, although upon induction (0.1 and 1.0 mM IPTG) a lower growth plateau and longer lag time were observed with respect to the empty plasmid control (Fig. 2A-B). This could be attributed to a metabolic burden from carrying the plasmid.

**Figure 2.**
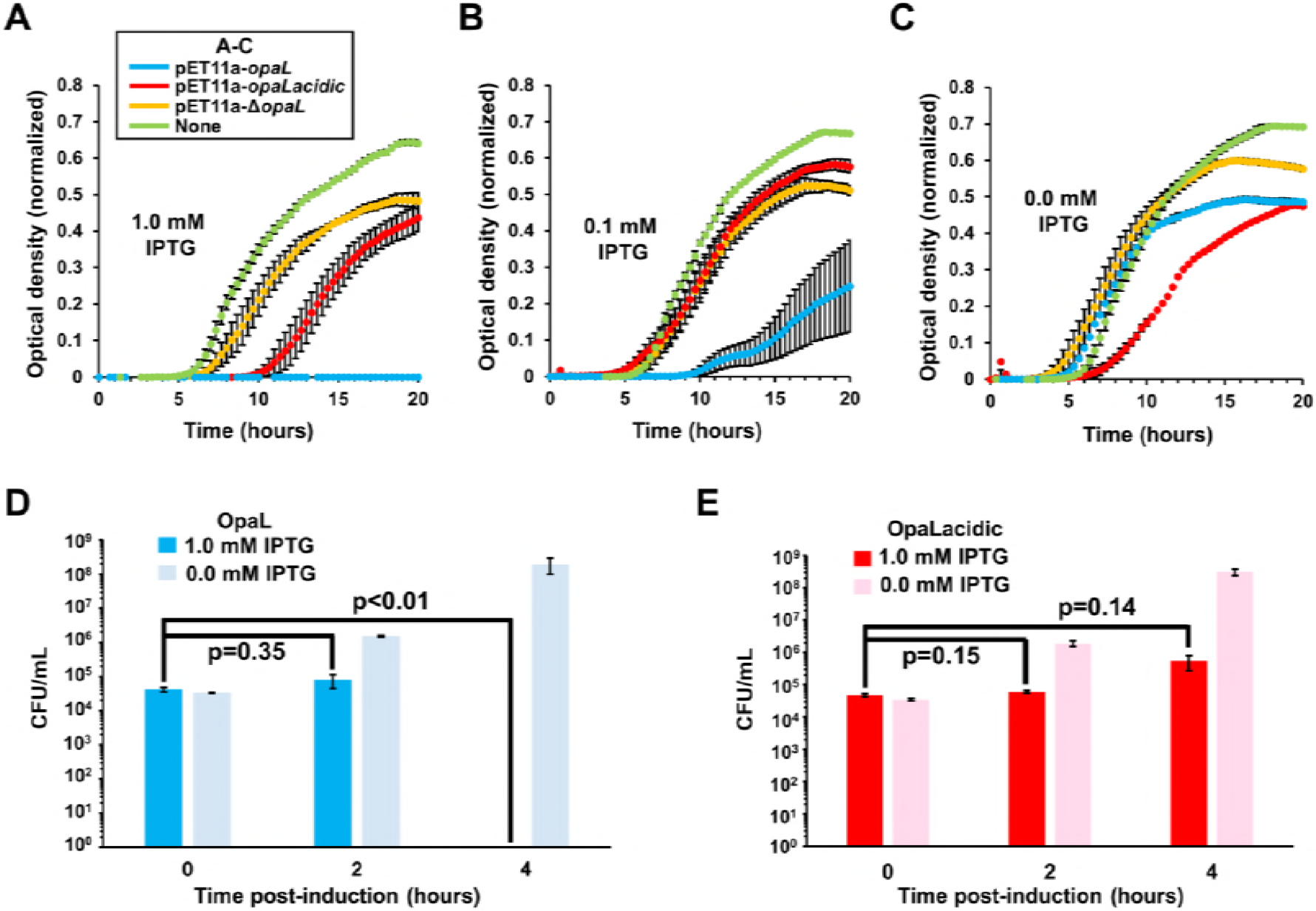
Antimicrobial effect of OpaL. **(A-C)** Growth curves of BL21 (DE3) carrying synthetic antimicrobial genes on plasmids pET11a-*opaL* (OpaL), pET11a-*opaLacidic* (OpaLacidic), pET11a-Δ*opaL* (control vector without an open reading frame), and no plasmid (None) under: **(A)** 1.0 mM IPTG induction, **(B)** 0.1 mM IPTG induction, and **(C)** without IPTG induction. **(D)** CFUs of BL21 (DE3) carrying pET11a-*opaL* and no plasmid at t=0, t=2, and t=4 hours post induction with 1.0 mM IPTG and at the same time points without IPTG. **(E)** CFUs of BL21 (DE3) carrying pET11a-*opaLacidic* and no plasmid at t=0, t=2, and t=4 hours post induction with 1.0 mM IPTG and at the same time points without IPTG. Data shown in this figure represent the means of three biological replicates. Error bars represent standard error and P-values were calculated using a two-tailed type II t-test.

In colony forming unit (CFU) experiments, the viable cell counts of *E. coli* BL21 (DE3) expressing OpaL dropped to zero after 4 hours of induction with 1.0 mM IPTG (p<0.01), indicating a bactericidal mechanism of action (Fig. 2D). In the absence of IPTG, the viable cell count continued to significantly increase over time. The viable cell counts after 2 hours and 4 hours for *E. coli* BL21 (DE3) carrying pET11a-*opaLacidic* were significantly lower with 1.0 mM IPTG than without any IPTG (Fig. 2E). Without IPTG, growth was observed at a decreased rate relative to the induced OpaLacidic cultures. This suggests that OpaLacidic likely operates by a bacteriostatic mechanism. Our data demonstrate that the OpaL and OpaLacidic both confer antimicrobial properties, but that OpaL has a much stronger effect.

### 3.3 OpaL causes protein aggregation

To test whether OpaL operates by an aggregative mechanism, we employed the statistical thermodynamics algorithm TANGO designed for prediction of aggregation-prone regions in proteins (18). We assumed OpaL and OpaLacidic concentrations of 2 mM based on the T7 promoter’s known expression levels (30). Using standard physiological parameters for *E. coli* (31, 32), we entered an ionic strength of 0.25 mM, a cytosolic pH of 7.5, and a temperature of 37°C. Across all its residues, OpaL showed a very high mean aggregation propensity of 35.7% (values greater than 5% are predictive of aggregation) (18), while OpaLacidic had a quite low mean aggregation propensity of 0.7% (Fig. 3A, supplementary table S2). These data support the idea that OpaL expression triggers the formation of lipophilic aggregates.

**Figure 3.**
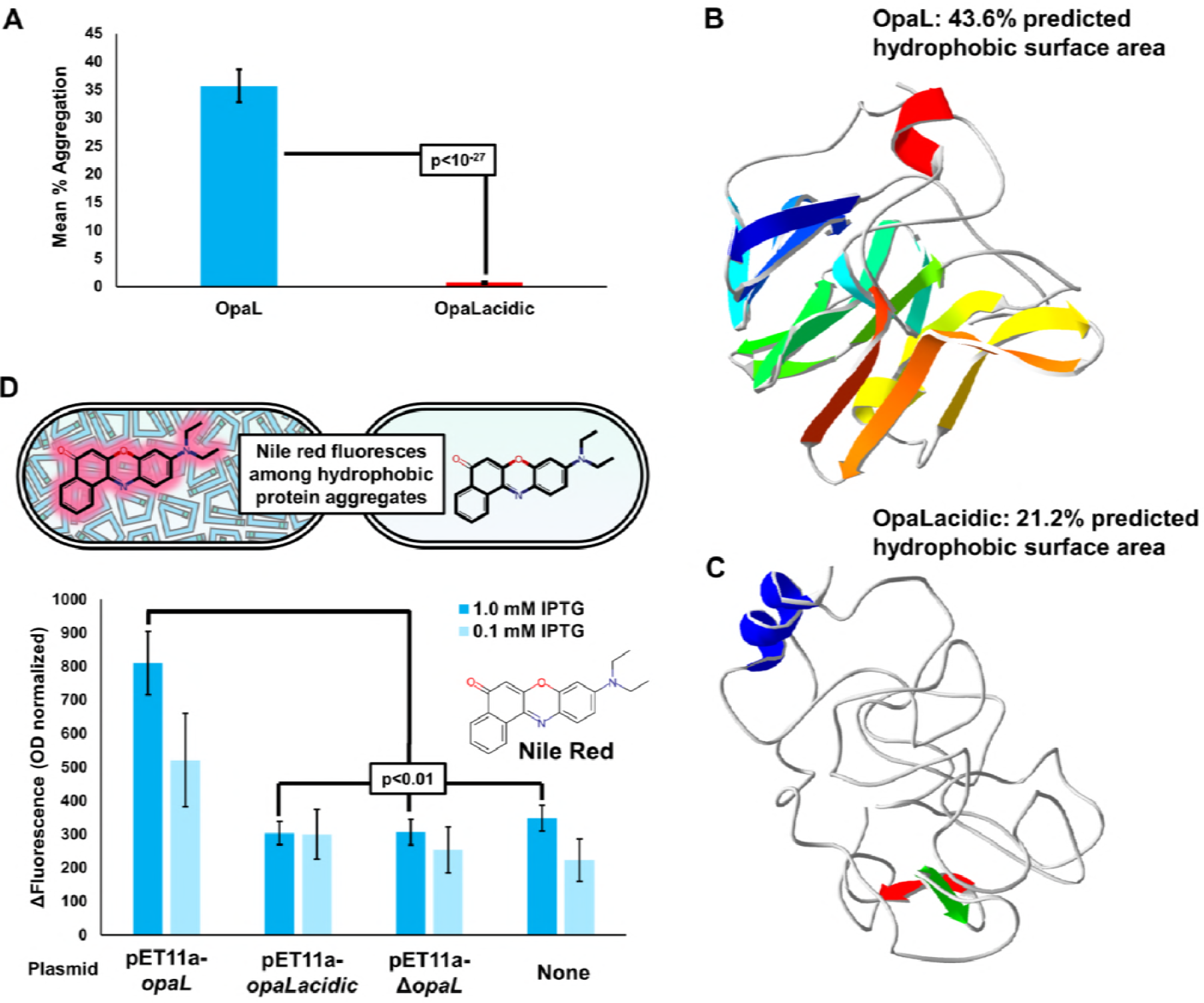
OpaL causes intracellular aggregate formation. **(A)** Predicted mean aggregation propensity percentages across all amino acids for OpaL and OpaLacidic using the TANGO statistical thermodynamics algorithm. **(B)** OpaL’s structure as predicted by the ab initio QUARK algorithm. This structure includes numerous β-sheets, similar to many naturally-occurring proteins which form aggregates (17). **(C)** The structure of OpaLacidic as predicted by the ab initio QUARK algorithm. This structure is mostly composed of unstructured loops, probably as a result of the 30 aspartic acid residues spaced regularly through the sequence. **(D)** BL21 (DE3) carrying pET11a-*opaL*, pET11a-*opaLacidic*, pET11a-Δ*opaL*, and BL21 (DE3) without any plasmids were stained with nile red to demonstrate that OpaL forms nonpolar aggregates. Fluorescence was measured immediately prior to IPTG induction and 2 hours after induction. These data represent the means of five biological replicates. In this figure, error bars represent standard error and *P*-values were calculated using a two-tailed type II t-test.

The QUARK algorithm, an ab initio protein structure prediction tool, was used to predict the tertiary structural characteristics of OpaL and OpaLacidic. DeepView was employed to visualize these structures and to compute their solvent-exposed hydrophobic surface areas. OpaL was predicted to be rich in β-sheet structures (Fig. 3B). By contrast, OpaLacidic was almost entirely composed of unstructured loops (Fig. 3C). The higher antibacterial toxicity of OpaL relative to OpaLacidic is consistent with these results since many pathological protein aggregates are also rich in β-sheets (17, 18). OpaL’s predicted structure demonstrated 43.6% hydrophobic surface area, while OpaLacidic demonstrated a predicted hydrophobic surface area of 21.2% (Fig. 3B-C). The frequent acidic residues in OpaLacidic are likely the reason that OpaL has more than twice the predicted hydrophobic surface area compared to OpaLacidic. Once again, OpaL’s higher antibacterial toxicity relative to OpaLacidic is consistent with these results since protein aggregation is known to be heavily dependent on hydrophobicity (15, 16, 18).

To provide experimental evidence for OpaL’s aggregative mechanism, we performed an aggregation assay by staining host *E. coli* BL21 (DE3) with the dye nile red. This dye fluoresces upon exposure to the hydrophobic environments of protein aggregates (33). When induced with 1.0 mM IPTG, cells carrying pET11a-*opaL* showed significantly higher increases in fluorescence after induction relative to cells with pET11a-*opaLacidic*, pET11a-Δ*opaL* or cells without any plasmids (p<0.01) (Fig. 3D). When induced with 0.1 mM IPTG, cells expressing OpaL showed higher increases in fluorescence after induction relative to the other groups, though the differences were not statistically significant at this lower IPTG concentration. This result demonstrates consistency with the design behind our peptides since the hydrophobic regions of OpaLacidic are periodically interrupted by 30 aspartic acid residues, while OpaL’s hydrophobic regions are much more continuous with only 7 aspartic acids. The repulsive charge-charge interactions between the aspartic acid residues might be responsible for this lesser propensity to form large aggregates and induce nile red fluorescence, especially since nile red fluorescence is not known to decrease under low pH conditions (34). These results are consistent with the data generated by the TANGO algorithm, further supporting OpaL’s mechanism of hydrophobic aggregation.

### 3.4 Delivering *opaL* by repurposing the RK2 conjugative plasmid

To utilize antimicrobial peptides in a viable therapeutic approach, appropriate strategies to deliver them to targeted pathogens are required. We show that bacterial conjugation serves as a potential approach to achieve this. Previous work has demonstrated this viability by delivering antibacterial CRISPR systems (4, 35, 36) and toxic hyper-replicating plasmids (37, 38). Here we developed a bacterial conjugation based delivery system for the de novo artificial gene which encodes OpaL (Fig. 4A). Unlike other antimicrobial peptides, the aggregate forming OpaL peptide is expressed intracellularly and in a genetically targeted fashion from its recipient-specific promoter. The *opaL* gene was placed under the control of this recipient-specific promoter, so only the recipient strain (not the donor) could transcriptionally activate *opaL.* Therefore, donor bacteria were unaffected while recipient growth inhibition occurred upon intracellular expression. By using the self-transmissible RK2 plasmid and a pET11a-*opaL* shuttle plasmid, *opaL* may exponentially spread through bacterial populations.

**Figure 4.**
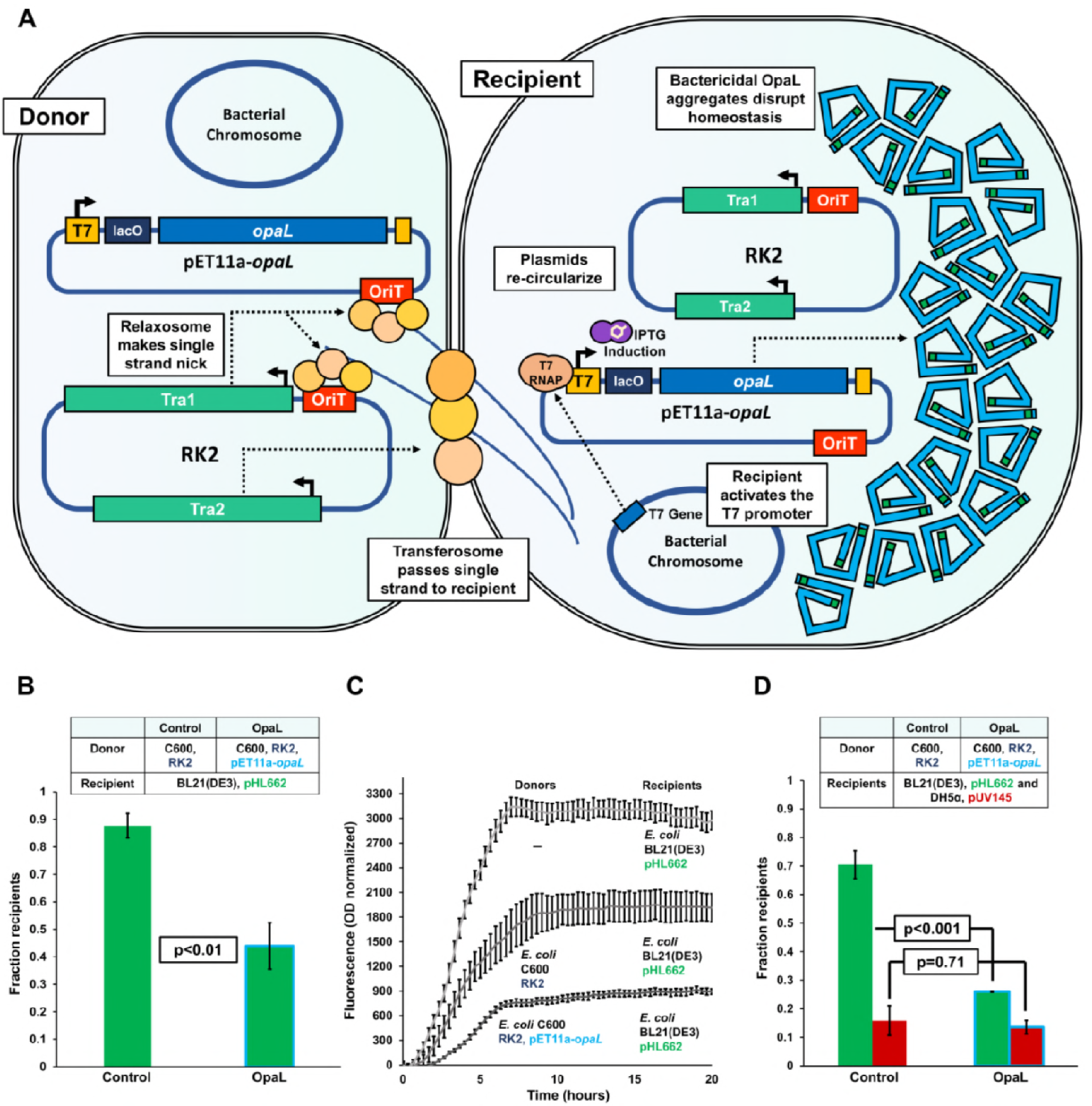
OpaL shows targeted killing when delivered by bacterial conjugation. **(A)** OpaL delivery using bacterial conjugation. Donor bacteria transfer the broad-host-range conjugative plasmid RK2 and the shuttle plasmid pET11a-*opaL* to recipient bacteria. The RK2 plasmid’s Tral operon encodes a relaxosome complex which makes a single-stranded nick in the origin of transfer (OriT) (47). RK2’s Tra2 operon encodes a transferosome complex which facilitates the transfer of the ssDNA to the recipient. The 450 bp OriT sequence from RK2 was cloned into pET11a-*opaL* so that *opaL* would undergo conjugative transfer. Using this mechanism, these conjugative vectors move into the recipient bacteria, re-circularize, and begin replicating (44). *E. coli* BL21 (DE3) encodes T7 RNA polymerase which binds the T7 promoter upstream of *opaL* and initiates expression. OpaL accumulates and forms insoluble aggregates, killing the host bacterium. **(B)** Two-strain CFU mating-toxicity assay. In the control co-cultures, donor *E. coli* C600 (RK2) and recipient *E. coli* BL21 (DE3) (pHL662) were used. In the experimental cocultures, donor *E. coli* C600 (RK2, pET11a-*opaL*) and recipient *E. coli* BL21 (DE3) (pHL662) were used. **(C)** Growth curves for mating-toxicity experiments. These growth curves include fluorescence (normalized to OD) for co-cultures of donor *E. coli* C600 (RK2) and recipient *E. coli* BL21 (DE3) (pHL662), co-cultures of donor *E. coli* C600 (RK2, pET11a-*opaL*) and recipient *E. coli* BL21 (DE3) (pHL662), and recipient mono-cultures of *E. coli* BL21 (DE3) (pHL662). These cultures were inoculated into a microplate after mating on solid medium for 5 hours. Recipient growth was measured by GFP fluorescence. These data represent the means of four biological replicates and error bars represent standard error. **(D)** Three-strain CFU mating-toxicity assay. In the control co-cultures, donor *E. coli* C600 (RK2), recipient *E. coli* BL21 (DE3) (pUV145), and recipient *E. coli* BL21 (DE3) (pHL662) were used. In the experimental co-cultures, donor *E. coli* C600 (RK2, pET11a-*opaL*), recipient *E. coli* BL21 (DE3) (pUV145), and recipient *E. coli* BL21 (DE3) (pHL662) were used. For panels B and D the mean recipient fractions relative to total cells after 5 hours of mating are displayed. These data represent the means of three biological replicates. Error bars represent standard error and *P*-values were calculated using a two-tailed type II t-test.

We chose the broad host range conjugative RK2 plasmid to facilitate delivery of the *opaL* gene because of RK2’s high transfer frequency, conjugative promiscuity, and stability (39, 40). RK2 was originally isolated from antibiotic resistant *Pseudomonas aeruginosa* and *Enterobacter aerogenes* strains at the Birmingham Accident Hospital in 1969 (40). Under optimal conditions, RK2, and its shuttle plasmids have very high conjugation frequencies (41–43). For instance, RK2 has been shown to mobilize shuttle plasmids from donor to recipient *E. coli* and from donor *E. coli* to recipient *P. aeruginosa* with conjugation frequencies of 8 and 0.2 transconjugants per donor (respectively) (43). The RK2 plasmid can be transferred among most gram-negative and many gram-positive bacteria and has even been shown to move from prokaryotic to eukaryotic cells, albeit at a much lower frequency (39, 41). Additionally, RK2 is known to be a stable plasmid as result of active partitioning, post-segregational killing, and multimer resolution (44). Active partitioning involves NTPases which transport copies of the plasmid to opposite poles of the cell prior to division (45). Post-segregational killing utilizes a slowly degraded toxin and rapidly degraded antidote. If the antidote’s production ceases as a result of plasmid loss, the toxin remains in the cell for longer than the residual antidote and the cell loses viability (46). In multimer resolution, resolvases correct plasmid multimers formed by recombination (44). These characteristics allow RK2 to rapidly spread plasmid DNA through bacterial populations. RK2 encodes relaxosome and transferosome proteins to mediate conjugative transfer (Fig. 4A). The relaxosome binds RK2’s origin of transfer (OriT) site and makes a single-stranded nick to relieve the plasmid’s negative supercoiling (47). The nicked DNA strand unwinds and moves to the membrane-bound mating transferosome complex which then transfers the plasmid to the recipient. In the recipient, the linear ssDNA recircularizes and the other strand is synthesized by rolling-circle replication (44).

To deliver *opaL* to recipient cells, we designed the pET11a-*opaL* shuttle plasmid, which includes a pET11a backbone, *opaL* cloned downstream of the T7 promoter and lac operator, a 450 bp sequence identical to the OriT site in RK2 (48), and a chloramphenicol (Cm) resistance gene (supplementary Fig. S1). The pET11a backbone was chosen over RK2 itself so that its higher copy number of 15-20 copies per cell (49) (with its pBR322 OriR) relative to RK2’s 4-7 copies per cell (50) and strong T7 promoter would maximize OpaL expression in targeted bacteria. Also, the much smaller (7 kb) pET11a-*opaL* plasmid enables more efficient genetic manipulation than would be possible if the large (60 kb) (48) RK2 plasmid had served as the backbone. Since the T7 promoter’s expression requires host cells with cytosolic T7 RNA polymerase, pET11a-*opaL* achieves strain-specific killing of *E. coli* BL21 (DE3), leaving host cells which lack the T7 RNA polymerase unharmed. Future extensions of this system may allow promoter-based targeting of pathogenic microorganisms, avoiding the killing of indigenous human microflora which occurs with most traditional antibiotics (51).

We demonstrated the conjugative transfer of RK2 by mating donor *E. coli* C600 carrying the RK2 plasmid with recipient *E. coli* XL1 Blue. To verify that pET11a-*opaL* could undergo mobilization by RK2, we performed the same procedure using *E. coli* C600 carrying both RK2 and pET11a-*opaL* as donors. After the mating these strains, we plated the mating cultures on X-gal and ampicillin (for RK2 matings) or Cm (for pET11a-*opaL* matings) in order to obtain CFUs of donors and transconjugants. We also plated the same cultures on X-gal without either antibiotic in order to obtain total recipient CFUs. Since *E. coli* XL1 Blue has the ΔlacZ mutation, transconjugant and recipient colonies were white, while donor colonies were blue. The conjugation frequencies were determined by calculating the ratio of transconjugant to total recipient colonies. The pET11a-*opaL* vector, when mobilized by RK2, showed a mean conjugation frequency of 2.63×10^−2^±1.23×10^−2^ transconjugants per recipient, confirming that RK2 is capable of conjugatively transferring pET11a-*opaL*.

### 3.5 OpaL kills recipient cells in the two-strain mating experiment

We mated donor *E. coli* C600 carrying RK2 and pET11a-*opaL* plasmids with recipient *E. coli* BL21 (DE3) carrying the pHL662 plasmid (which expresses GFP) using 1:3 donor to recipient ratios based on optical density in presence of 1.0 mM IPTG. As a control, we also performed these matings using donors that carried only the RK2 plasmid under similar conditions. After the matings had been completed, the cultures were diluted and spread on fresh solid medium containing 1.0 mM IPTG. Using fluorescent protein expression to distinguish between strains (supplementary Fig. S2A), we calculated the ratios of recipient colonies to total colonies. The experimental group’s recipient CFU fractions were 2.0-fold lower than those in the mating control (p<0.01), demonstrating successful *opaL* delivery and toxicity (Fig. 4B). In addition to the CFU experiments, we measured recipient growth curves using GFP fluorescence from mating cultures in a microplate reader. The recipient *E. coli* BL21 (DE3) showed significantly decreased growth when donors delivered both RK2 and pET11a-*opaL* as compared to the control without pET11a-*opaL* (Fig. 4C). After 20 hours of growth, the mean fluorescence value of the experimental group was 1.5-fold lower than that the control (p<0.001).

### 3.6 Strain-specific killing in a three-strain mating experiment

To show this proof-of-concept in a more complex competitive environment, we next conducted three-strain matings between donor *E. coli* C600 (carrying RK2 in the control, with both RK2 and pET11a-*opaL* for the experimental group), targeted recipient *E. coli* BL21 (DE3) with GFP expressed from pHL662, and a non-targeted recipient *E. coli* DH5α with mCherry expressed from the plasmid pUV145 (22) (supplementary Fig. S2B). The OD-adjusted ratio of donors to targeted recipients to non-targeted recipients was 1:1:2. Otherwise, the same general procedures used in the two-strain matings were employed. Both the fractions of targeted recipients and the fractions of non-targeted recipients were determined relative to the total colonies on each plate. The experimental group’s targeted recipient CFU fractions were a significant 2.7-fold lower than those in the mating control (p<0.001), while the non-targeted recipient CFU fractions did not differ significantly from the control (p=0.71) (Fig. 4D). This demonstrates that our system may operate effectively with more complicated competition dynamics and that non-target recipient bacteria are not harmed by *opaL*.

## 4 Discussion

We designed and tested the first aggregating de novo antimicrobial peptide, OpaL, to form insoluble inclusion bodies and disrupt bacterial homeostasis, serving as the basis for the rational design of novel antimicrobials. We envision that OpaL resistant strains may arise less rapidly than small-molecule antibiotic-resistant strains since hydrophobic aggregates do not bind to a specific target and hydrophobic aggregates disrupt homeostasis through numerous mechanisms (15, 16). Therefore, the target site mutations implicated in typical resistant phenotypes (52) may be probabilistically less likely. It is also unlikely that insoluble protein aggregates would be removed via efflux mechanisms. Furthermore, the peptide’s de novo amino acid sequence could decrease the frequency of enzymatic exaptation towards specific binding and cleaving of motifs within OpaL. It should be noted that mutations in the promoter upstream of *opaL* could decrease intracellular OpaL accumulation, providing a route for outgrowth. In this scenario, donors with fresh copies of the original *opaL* gene could be supplied to restore full expression. Even if resistance was to arise, OpaL’s design is highly amenable to directed evolution. Since OpaL’s mechanism depends on persistent “misfolding” and aggregation rather than fixed tertiary folds, mutations should be less likely to decrease OpaL’s activity, widening the pool of potentially improved mutants as compared to most protein therapeutics (53). After iterated mutagenesis and screening, this scenario’s *opaL* gene may regain activity against resistant pathogens. Furthermore, as hydrophobic aggregation can occur in any aqueous cellular environment, OpaL’s mechanism may enable activity in diverse types of targeted bacteria.

We also demonstrate an assembly of biological components to develop a mobile genetic system which eliminates promoter-targeted strains within bacterial populations. We employed the highly efficient RK2 plasmid to mobilize our pET11a-*opaL* shuttle vector and spread *opaL* through populations of bacteria. Our RK2 mediated delivery system has promise for treating infections which involve biofilms. The rate of bacterial conjugation tends to be much higher in biofilms (which pathogenic bacteria such as *E. faecalis* and *P. aeruginosa* frequently form) (54, 55) than in laboratory culture conditions. Of particular note, Hausner and Wuertz showed that conjugative transfer increases up to 1,000-fold when using RK2 derived plasmids in biofilms (56). In large part, this occurs because cells within biofilms are immobilized and consequently form more stable conjugative junctions (57). We envision pathogenic biofilms providing a rich conjugative environment for *opaL* propagation. Donor bacteria with similar surface characteristics to the target pathogens might be capable of integrating into biofilms. OpaL’s effectiveness could be tested in the future using pathogenic bacteria such as *P. aeruginosa* under biofilm forming conditions. Our de novo design approach for an aggregating antimicrobial peptide may be adaptable to many types of infections, yet possess specificity in its targeting of pathogens. This technology provides new opportunities for addressing the global challenge of antibiotic-resistant bacterial infections.

## ACKNOWLEDGEMENTS

This work has been supported by William M. Keck Foundation and National Science Foundation award number MCB1714564 to AC. PO and CMC were supported by National Science Foundation Fellowships. The funders had no role in study design, data collection and analysis, decision to publish, or preparation of the manuscript.

## AUTHOR CONTRIBUTIONS

AC supervised the project. LC, PO, CMC and AC designed the research. LC performed the experiments and the statistical data analysis. LC, AC, PO and CMC wrote the manuscript.

The authors report no conflict of interest.

